# Protein-ligand co-design: a case for improving binding affinity between Type II NADH:quinone oxidoreductase and quinones

**DOI:** 10.1101/2024.06.11.598532

**Authors:** Vladimir Porokhin, Anne M. Brown, Soha Hassoun

## Abstract

Biological engineering aims to enhance biological systems by designing proteins with improved catalytic properties or ligands with enhanced function. Typically, applications permit designing proteins, e.g., an enzyme in a biodegradation reaction, or ligands e.g., a drug for a target receptor, but not both. Yet, some applications can benefit from a more flexible approach where both the protein and ligand can be designed or modified together to enhance a desired property. To meet the need for this co-design capability, we introduce a novel co-design paradigm and demonstrate its application to Ndh2-quinone pairings to enhance their binding affinity. Ndh2, type-II NADH dehydrogenase, is an enzyme found in certain bacteria that facilities extracellular electron transfer (EET) when interacting with exogenous quinone mediators. This interaction leads to the generation of a detectable electric current that can be used for biosensing applications. Our results demonstrate the benefits of the co-design paradigm in realizing Ndh2-quinone pairings with enhanced binding affinities, therefore highlighting the importance of considering protein-ligand engineering from a holistic co-design perspective.

## 1 Introduction

Synthetic biology seeks to redesign biological systems for performing specific tasks by engineering their constituents, such as protein sequences and molecules. Protein engineering aims to identify mutants with improved properties, such as selectivity, specificity, expression, solubility, and stability. As the combinatorial mutational space of protein sequences is extremely large, a purposeful approach is required to design and explore protein variants. A popular method for protein engineering is directed evolution (DE), where proteins undergo iterative rounds of gene diversification and screening, mirroring the process of natural evolution on an accelerated timescale. Although labor-intensive [1], this method has been successfully used for evolving biological pathways and improving catalytic properties of enzymes [1]. Complementing the purely experimental approach of DE, rational design [2, 3] leverages structural and mechanistic properties of proteins to alter function in a desirable way. In addition, a growing selection of computational methods is available, from heuristic-based optimization techniques [4, 5], to highly parallelized molecular dynamics [6] and machine learning (ML) models [7, 8, 9], helping to elucidate protein-function relationships in a less expensive fashion. Ligand design aims to select molecules with specific properties, such as affinity, druglikeness, metabolic stability, and bioavailability [10]. However, the space of molecular structures is also extremely large: for example, the number of pharmacologically relevant structures alone may exceed 10^60^ [11]. To address this challenge, computational approaches have emerged, spanning fragment-based ligand design to evolutionary algorithms [12]. Several ML methods have also been introduced [13, 11].

A common modality of existing design methods is their assumption of the *individual design* paradigm, where proteins and ligands are designed individually. Individual design is appropriate in situations where either the protein or the ligand is allowed to vary and the other is fixed, for example when developing a drug to target a particular protein or when an enzyme is mutated to maximize yield of a particular product. However, there are emerging applications, most notably in bioelectronic sensing [14] and synthetic biosynthesis [15], where it is desirable to engineer *both* the ligand and protein. A *co-design* paradigm would allow for simultaneous changes of ligand and protein, and the consideration of how the change of one component affects the function of the other or the interaction between them. For example, binding affinity is dependent on both enzyme and ligand, and in the space of possible co-modification, more favorable pairings may be identified by matching the binding pocket volume to the size of the molecule and vice versa.

One example of an application that benefits from co-design is the case of type II NADH:quinone oxidoreductase, also known as Ndh2, and its interaction with quinones. Ndh2s comprise a family of membrane-bound proteins acting on nicotinamide adenine dinucleotide (NAD), responsible for oxidation of NADH and reduction of quniones. They play a central role in respiratory chains of many organisms [16] and help maintain the balance of NADH and NAD^+^ [17]. Ndh2 has been proposed as a drug target for treatment of numerous infectious and chronic diseases [16, 18, 17, 19], and like other oxidoreductases, may be of interest in measurement and pollution control applications [20]. Recently, an extracellular electron transfer (EET) pathway has been described in *Lactiplantibacillus plantarum* that can reduce an extracellular electrode in the presence of exogenous quinones using Ndh2 as a mediator [21]. The rate of EET varied across different quinone analogs and concentrations, creating distinguishable electronic signals and presenting an exciting opportunity for building inexpensive whole-cell bioelectronic sensors for pharmacologically relevant quinones. However, improving sensitivity or selectivity of this sensing system through Ndh2 and/or quinone co-design remains a challenge, with only a small number of quinones currently characterized for this purpose [22, 23].

We investigate in this work a method for the computational co-design of protein-ligand pairs and present its application to Ndh2-quinone binding for EET rate maximization. Our proposed paradigm, referred to as “co-design,” aims to simultaneously explore the space of ligand and protein modifications that can lead to improved system design. In the case of EET, we seek to improve binding affinity over the Ndh2-quinone complexes currently known for facilitating EET. Our goal herein is therefore to show that the co-design paradigm can yield enhanced biological systems over individual design paradigms.

Our approach consists of three major steps. First, we build libraries of protein and quinone variants by considering changes likely to impact their mutual binding. For protein variants, we identify a number of residues in close proximity to the quinone binding site and enumerate all possible single amino acid substitutions in those locations. For ligand variants, we construct modified quinones by iteratively adding functional motifs such as phenyl and hydroxy groups to one of the two EET-active molecules: 1,4-dihydroxy-2-naphthoic acid (DHNA) and menadione [22]. Second, we implement two strategies to search the protein-ligand space for pairings with improved binding affinity: individual design and co-design. In individual design, we explore protein-ligand pairs where only the protein or only the ligand has been modified, as is common in current design paradigms. Meanwhile, with co-design, we consider all combinations of proteins and ligands in our libraries. Third, we evaluate the resulting combinations using molecular docking with AutoDock Vina [24]. The binding affinity predicted by AutoDock Vina represents the favorability of protein-ligand interaction, which we believe is a key contributor to EET activity. However, limited receptor flexibility is one of the major shortcomings of molecular docking methods [25] and is commonly addressed by leveraging the more accurate molecular dynamics simulations [26]. As such, we subsequently validate our proposed combinations using predicted Molecular Mechanics with Generalized Born and Surface Area Solvation (MM/GBSA) based free energy of binding calculations from Schrödinger Maestro [27, 28, 29]. Our results show that the co-design paradigm is more comprehensive than individual design as it considers a wider range of proteins-ligand combinations. Further, we demonstrate that the pairings identified by co-design yield more favorable interactions when compared to pairings identified through individual design. We also present a selection of ten Ndh2-quinone pairs identified by our method and describe their key features relevant to EET.

## 2 Methods

### 2.1 Modeling *L. plantarum* Ndh2

The sequence of the *L. plantarum* Ndh2 protein was obtained from the UniProtKB database [30] (entry D7VAS3). Structural homology models were then constructed using AlphaFold [31, 32] and Robetta [33] (RoseTTAFold [34] model) web servers, two state-of-the-art protein structure prediction methods. To ensure consistent positioning of residues, cofactors, and ligands across the structures, one of the Robetta models was arbitrarily chosen as the coordinate reference and all other models were oriented to it using the PyMOL Alignment [35] plugin.

The crystal structure (PDB ID 4G73) [36] of an Ndh2 variant sourced from another organism, *S. cerevisiae* S288C, was used to validate and augment the homology models. This structure from Feng et al. [36] is one of the few depictions of an Ndh2-quinone complex we could locate in the Protein Data Bank (PDB). Feng et al. also proposed an alternative (PDB ID 4G74) of the Ndh2-quinone complex; however, it included a Triton X-100 molecule, which would not be present *in vivo*, so we removed this molecule. The chosen structure from *S. cerevisiae* S288C had 26% sequence identity to the *L. plantarum* Ndh2 as measured by blastp [37] and presented a similar structure, yielding alignment RMSDs of 1.295 Å and 1.457 Å for AlphaFold and Robetta models, respectively.

The *S. cerevisiae* S288C crystal structure also included poses for flavin adenine dinucleotide (FAD), NADH, and a quinone (Ubiquinone 25) bound to the protein. Those poses were overlaid on our homology models, providing the locations of the cofactors as well as the binding pocket for the quinone. The presence of NADH is not required under all proposed Ndh2 catalysis mechanisms. In particular, the ping-pong mechanism assumes that NADH and quinone are not bound to the protein at the same time, but react with it sequentially [16]. As such, we excluded the NADH pose from our subsequent modeling, but retained FAD. The models were evaluated based on their ability to dock with the quinone and their overall structure quality, and the AlphaFold model was selected moving forward; see the “Results and discussion” section for more details.

### 2.2 Protein and ligand preparation

A library of Ndh2 mutants was constructed by enumerating all single-point mutations of residues comprising the quinone binding pocket. These residues were manually selected based on their close proximity to DHNA when docked with our model of Ndh2. Three contiguous ranges of amino-acid positions were chosen for mutation: 320-322, 353-355, and 381-390. The PyMOL Mutagenesis tool [35] was used to introduce the mutations. In instances where multiple side chain orientations were possible, we selected the most probable rotamer predicted by PyMOL.

The resulting mutants were further analyzed to consider the favorability of their corresponding protein mutations. The Schrödinger Maesto [27] Residue Scanning tool was used to predict the change in Prime MM/GBSA free energy (“Δ Prime” hereafter) caused by all single amino acid substitutions. A negative change describes a mutation more favorable than the wildtype, while a positive change indicates the opposite. Changes greater than +10 kcal/mol generally imply a major structural change in the protein, which would likely be detrimental to its ability to bind to a quinone and/or perform EET. Therefore, all mutated variants with Δ Prime exceeding that threshold were discarded from further consideration.

A library of ligands was constructed by iterative addition of physiologically relevant functional groups to base molecules: initially DHNA and menadione, the two quinones known to facilitate EET in *L. plantarum* Ndh2 [22]. The functional groups were attached to heavy atoms on the periphery of either DHNA or menadione, with the exception of the carbonyl oxygen atoms in positions 1 and 4 (see Figures 5e and 5f) as to not impede the quinone ⇋ semiquinone transformation required by EET [22]. The functional groups included benzene, C_2_H_5_, CF_3_, CH_2_OH, CH_3_, CHF_2_, F, NH_2_, OCH_3_, and OH. [38, 39, 40] In addition, we considered groups CH_2_NH_2_, NO_2_, and C_4_H_4_NH; however, these motifs did not yield any useful molecules in our process and were excluded accordingly. The addition of a functional group was implemented as a generic reaction described by a SMARTS pattern [41] and applied using RDKit [42]. The resulting derivative molecules were validated for basic chemical validity using RDKit and cross-referenced against PubChem [43]. Invalid molecules or those not found in the database were excluded from further consideration. This process was repeated up to 6 times to produce all variant molecules in the library.

### 2.3 Search strategy

We considered two strategies in this study: individual design and co-design. In individual design, only the protein or the ligand is being varied, with the other assumed as unchanged. Therefore, to generate protein-ligand pairs, we exhaustively enumerated all combinations featuring either (1) the unmodified DHNA and menadione quinones and one of the Ndh2 variants from our mutant library, or (2) the wildtype Ndh2 protein and a molecule from our ligand library. The former set corresponded to design options of the protein, while the latter provide design options for the ligand.

In co-design, modified proteins and ligands are being considered for evaluation. This strategy can identify pairs featuring both a mutated Ndh2 vairant and a modified quinone molecule. Unfortunately, the combinatorial space of such pairs is large, so the exhaustive enumeration approach we used for individual design would not be feasible. Instead, we randomly sampled a number of pairs corresponding to a subset of all possible protein-ligand pairs accessible to co-design.

### 2.4 Molecular Docking

The binding of *L. plantarum* Ndh2 variants and ligands was modeled using AutoDock Vina [24] with bounding box dimensions 15Å x 17Å x 20Å and center position (38, 44, 2), referenced to the Robetta model selected. The size and placement of the box were manually chosen as to include the quinone binding pocket while rejecting most of the poses inconsistent with the expected quinone orientation with respect to FAD and the protein. The exhaustiveness of the search was set to 8, the maximum number of binding modes to be generated was 9, and the maximum energy difference between displayed modes was set to 3 kcal/mol. When multiple poses were possible, we selected the one with the most favorable predicted binding affinity.

### 2.5 Evaluating protein-ligand pairings

The obtained pairings were evaluated based on the binding affinity predicted by AutoDock Vina as a part of molecular docking, as well as the predicted free energy of binding calculated using MM/GBSA. To run MM/GBSA, first each protein-ligand pair was converted into the compressed Maesto (”maegz”) format. The Ndh2 variant structure was refined with the Protein Preparation Wizard (prepwizard) [44]. The quinone was prepared using the LigPrep program [45]; since this operation moved the molecule away from its docked pose, the LigPrep-processed ligand was subsequently re-aligned to the original output from AutoDock Vina. Finally, the prepared protein and quinone were combined into one complex and forwarded to the Prime program [28, 29] for the MM/GBSA calculation. This workflow was automated using the JobDJ toolkit and command line utilities available in the Schrödinger Maesto suite [27].

Protein-ligand combinations were filtered by MM/GBSA free energy of binding values, removing ones with positive predicted free energy of binding, as this was indicative of energetically unfavorable pairings. Finally, the remaining pairings were sorted by their AutoDock Vina binding affinities, and the top 10 were chosen for in-depth consideration.

## 3 Results and discussion

### 3.1 Structural model assessment

The modeled structures of the *L. plantarum* Ndh2 protein were first validated by ensuring they can dock with DHNA and menadione, the two quinones known to interact with the protein. Although both AlphaFold and Robetta homology models were considered, only the former yielded meaningful poses. Upon closer inspection of the residues in the pocket, we identified an unfavorable orientation of the Y390 side chain, exclusive to the Robetta models. Such orientation resulted in a clash with the expected position of the quinone that was implied by the placement of Ubiquinone-25 in the reference *S. cerevisiae* S288C Ndh2 structure (Figure 1). Replacement of Y390 with alanine resolved the issue, confirming the unfortunate orientation of the side chain as the cause. However, because the original Robetta structure wasn’t effective in modeling quinone binding, we chose to proceed with the AlphaFold structure from this point on.

**Figure 1:**
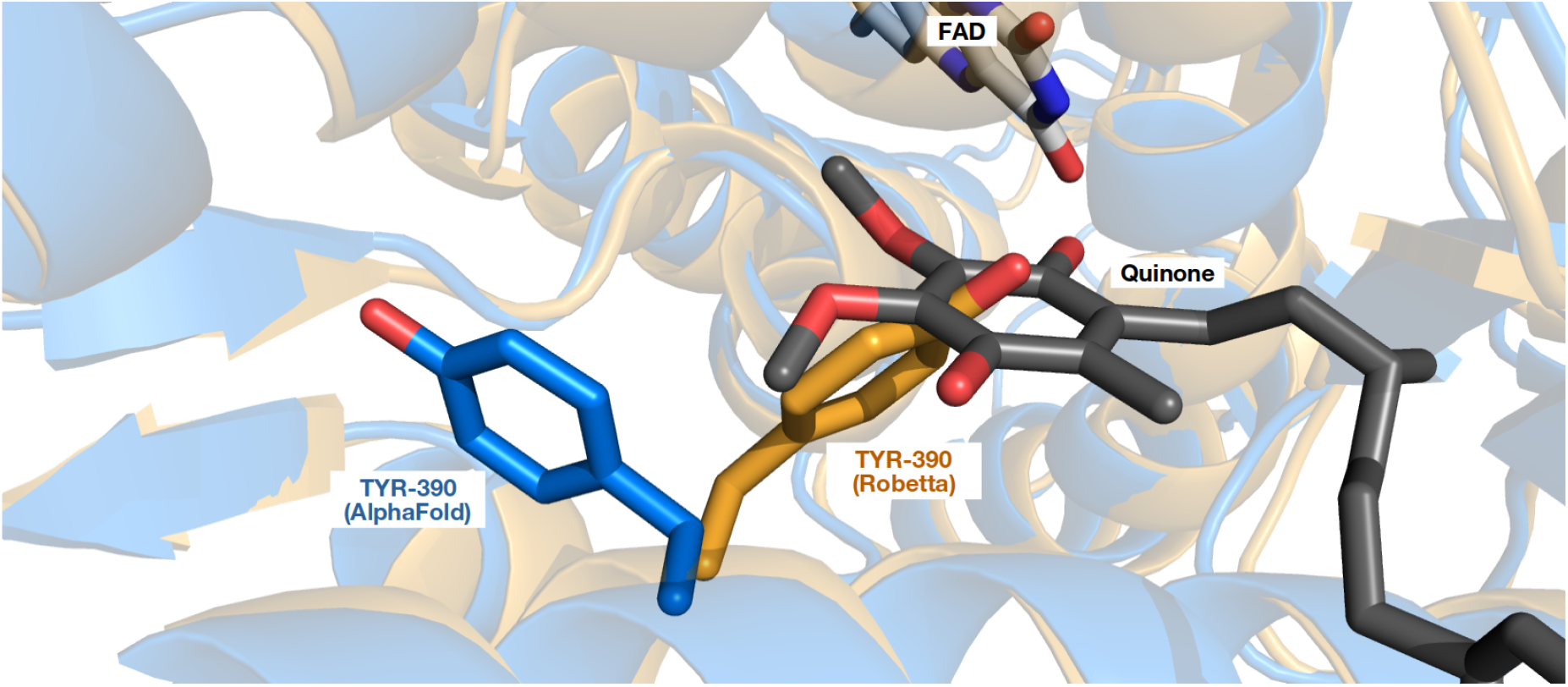
Y390 residue placement in AlphaFold (blue) and Robetta (orange) models. The expected position of the quinone (dark gray), superimposed from the *S. cerevisiae* S288C Ndh2 structure, demonstrates a clash unique to the Robetta model.

The selected homology model was then analyzed for quality using the SWISS-MODEL [46, 47, 48, 49, 50] and ProSA [51, 52] web servers, which indicated that our model had favorable structural characteristics in terms of (Φ, Ψ) residue angles, Z-score, and QMEAN assessment as shown in Figure S1.

To explore the characteristics of the *L. plantarum* Ndh2, we compared the residues in its quinone binding pocket with those in the *S. cerevisiae* variant (see Figure 2). Although several residues have mutual counterparts in both proteins and enclose the quinone in a similar manner, there are a number of differences that may contribute to unique behavior of the *L. plantarum* Ndh2. For example, the Q353 residue corresponding to L444 in 4G73 [36] has a different affinity to water, which may significantly alter the interaction with the quinone given the pocket’s proximity to the cell membrane. Additionally, two residues, E324 (H397 in 4G73) [36] and K383 (Y482), differ in the presence of a positively charged side chain, which may effect the orientation of the ligand. Those differences emphasize the need for further study of the *L. plantarum* Ndh2; however, the ability to dock DHNA and menadione in a similar position to the previously reported crystal structure lends confidence in our predicted model.

**Figure 2:**
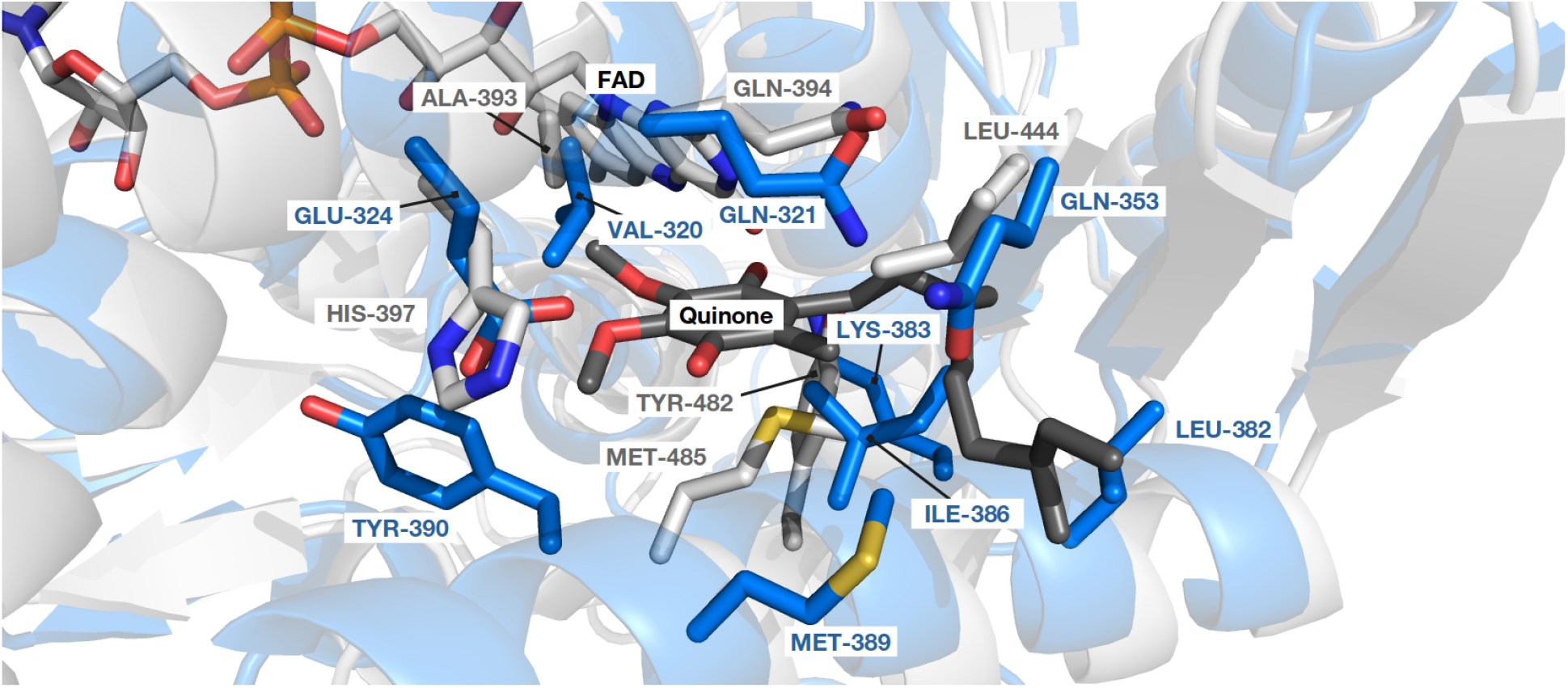
Overlay of *L. plantarum* (blue) and *S. cerevisiae* (light gray) structures shows the common features of the quinone (dark gray) binding pocket as well as the differences unique to each variant. Labels indicate the residues of interest.

### 3.2 Co-design allows richer design options

By allowing variability in both the protein and the quinone, the co-design approach considers a far greater number of possible combinations. In particular, when individual design focused on the protein and the ligand was limited to 366 and 822 pairings, respectively, co-design of both could target in excess of 150,000 pairs using the same protein and molecule libraries. In this study, we employed a sampling strategy for co-design, retrieving just over 15,000 pairings. Although exploring the larger quantity of pairs increases computational costs, it provides more opportunities for identifying favorable combinations.

The co-designed protein-ligand combinations spanned a wide range of AutoDock Vina binding scores and outnumbered those evaluated by individual design for any given affinity. Across the entire range of observed binding affinities, 86% or more of the pairings were derived from co-design; ligand and protein individual design contributed only up to 6% or 11% of the pairs, respectively. Furthermore, the pairs with the most improved binding affinities were only attainable by the co-design approach and not individual design (Figure 3).

**Figure 3:**
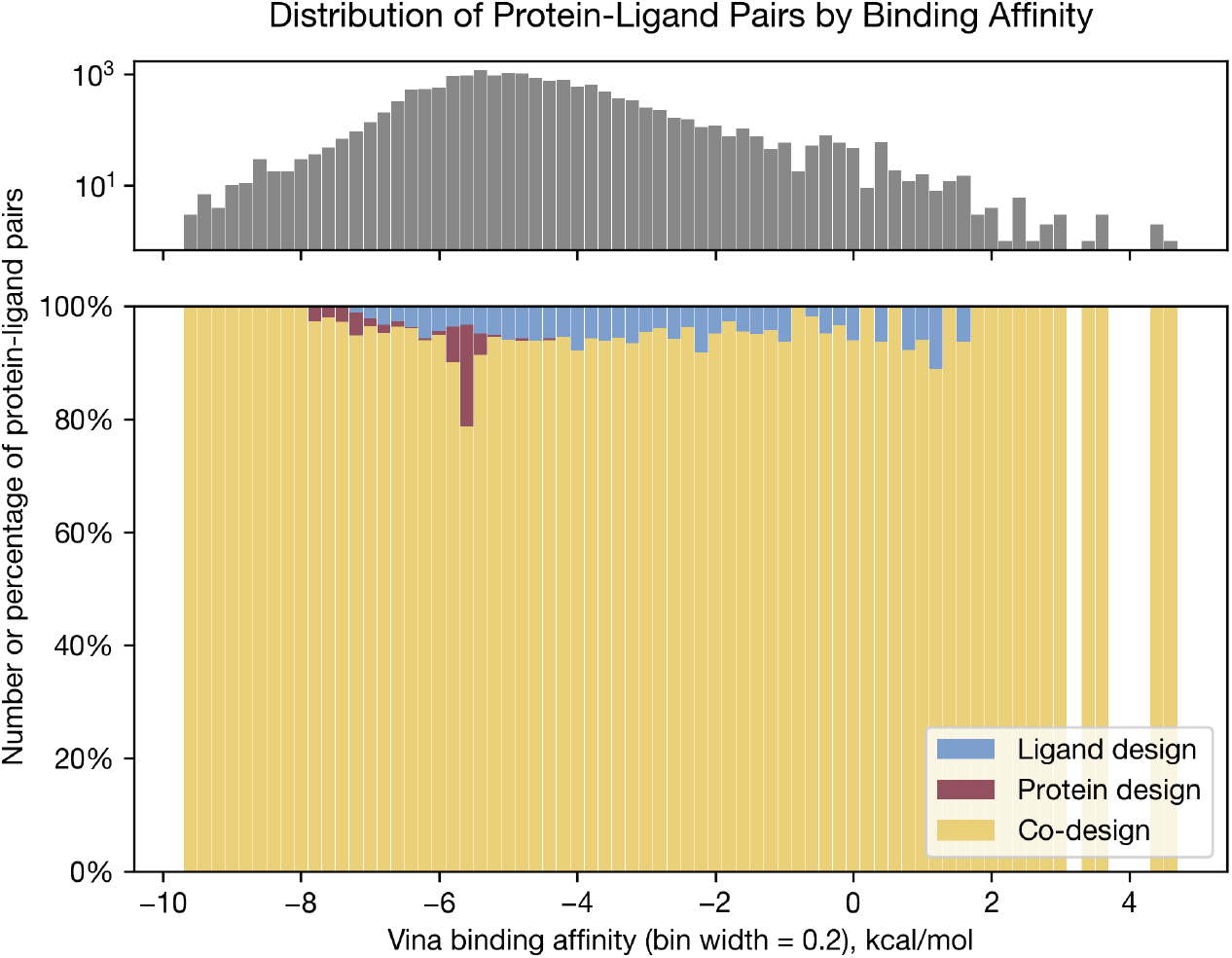
Distribution of protein-ligand pairs with respect to binding affinities. Top panel: overall number of pairs explored in this work. Bottom panel: relative percentage of protein-ligand pairs considered by each design strategy. The percentage of pairs evaluated by co-design (yellow) was significantly larger than that evaluated by ligand (blue) or protein (maroon) design, irrespective of the binding affinity, and the optimum affinities are only attainable via co-design.

It should be noted that both ligand and protein design strategies were exhaustively enumerated, while the co-design space was only partially sampled. As such, it is likely that co-design could achieve an even greater contribution, were it not subject to the limitations of sampling.

### 3.3 Co-design identifies more favorable pairings

Compared with individual design approaches, co-design identifies more favorable protein-ligand pairings. Overall, co-design explores a larger selection of options, which results in higher standard deviation and reduced average performance as measured by both AutoDock Vina binding affinity and MM/GBSA free energy of binding. However, when selecting for the top 50 pairs by AutoDock Vina binding affinity, this pattern reverses. For example, the average binding affinity over top 50 pairings was -6.2 (*±* 0.6) kcal/mol and -6.5 (*±* 0.2) kcal/mol for protein and ligand individual design, respectively. However, for co-design, the average AutoDock Vina binding affinity was more favorable, -8.9 (*±* 0.3) kcal/mol. This trend held for MM/GBSA free energy of binding values as well: the average free energy of binding over the same 50 pairings was -21.10 (*±* 7.69) kcal/mol and 9.31 (*±* 22.29) kcal/mol for protein and ligand design, respectively. Meanwhile, for co-design, the free energy of binding was once again more favorable, -34.83 (*±* 9.59) kcal/mol. Additional comparisons are provided in Table 1.

**Table 1:** Comparison of individual and co-design strategies. Average and standard deviation values are given for AutoDock Vina binding affinity and MM/GBSA free energy of binding; bolded text indicates the best value in each category. Average performance across all pairings is better for individual protein design. However, the performance over the top 50 pairings selected by AutoDock Vina is significantly better for the co-design approach.

| Metric | Design Approach | Top 50 pairs by AutoDock Vina | All pairings |
| --- | --- | --- | --- |
| AutoDock Vina binding affinity score, kcal/mol | Individual, protein | -6.2 ± 0.6 | **-5.6 ± 0.4** |
| | Individual, ligand | -6.5 ± 0.2 | -4.5 ± 1.3 |
| | Co-design | **-8.9 ± 0.3** | -4.6 ± 1.5 |
| MM/GBSA free energy of binding, kcal/mol | Individual, protein | -21.10 ± 7.69 | **-19.15 ± 11.65** |
| | Individual, ligand | 9.31 ± 22.29 | 8.92 ± 32.32 |
| | Co-design | **-34.83 ± 9.59** | 12.48 ± 61.50 |

For all but three protein variants, the co-design approach improves AutoDock Vina binding affinity over individual protein design. Across all mutation sites and replacement amino acids, co-design commonly achieved an improvement of over -1 kcal/mol, with a maximum of -2.3 kcal/mol. For the V320R, I386E, and Y390R mutants, regressions of +0.4, +0.2, and +0.2 kcal/mol, respectively, were observed in AutoDock Vina binding affinities. In these cases, the best pairing found by co-design had lower interaction affinity than the mutant-DHNA/menadione pair found by individual design. However, co-design could also propose the same pair, were it not limited by the number of random samples it was allowed to make. As such, this regression is not indicative of limitations of the co-design strategy. More details can be found in Figure S2.

### 3.4 Biochemical insights uncovered by co-design

The top 10 co-designed protein-ligand pairs ranged in binding affinities from -9.7 kcal/mol to -9.3 kcal/mol, which compared favorably to the best interaction affinities achieved by individual design approaches, -7.7 kcal/mol for protein design and -7.5 kcal/mol for ligand design. These pairings also had favorable MM/GBSA free energies of binding, ranging from -25.3 to -49.4 kcal/mol. The full list of the top 10 protein-ligand pairings is given in Table 2 and drawings of relevant ligands can be found in Figure S3.

**Table 2:** Favorable protein-ligand pairs suggested by the co-design approach. Ligands are identified by their IUPAC names as well as PubChem compound ids (indicated in parentheses). Binding affinity is reported by AutoDock Vina; free energy of binding is calculated using Prime MM/GBSA.

| Rank | Ligand | Protein variant | Pair type | Binding affinity, kcal/mol | Free energy of binding, kcal/mol |
| --- | --- | --- | --- | --- | --- |
| 1 | 2-Amino-3-(1-phenylethyl)naphthalene-1,4-dione (90135914) | I386D | 1 | -9.7 | -25.3 |
| 2 | 2-(1-Hydroxy-1-phenylethyl)naphthalene-1,4-dione (23730319) | I386C | 2 | -9.5 | -44.5 |
| 3 | 2-Amino-3-(1-phenylethyl)naphthalene-1,4-dione (90135914) | I386C | 1 | -9.5 | -36.1 |
| 4 | (2-Methylphenyl) 1,4-dihydroxy-5-methylnaphthalene-2-carboxylate (140286449) | I386S | 2 | -9.4 | -36.0 |
| 5 | (2-Methylphenyl) 1,4-dihydroxy-5-methylnaphthalene-2-carboxylate (140286449) | I386N | 2 | -9.4 | -29.5 |
| 6 | 2-(1-Hydroxy-3-phenylpropyl)naphthalene-1,4-dione (71652350) | I386A | 3 | -9.4 | -32.0 |
| 7 | 2-[Hydroxy(2-hydroxyphenyl)methyl]naphthalene-1,4-dione (245768) | I386S | 2 | -9.3 | -41.8 |
| 8 | (4-Hydroxyphenyl) 1,4-dihydroxy-3-methylnaphthalene-2-carboxylate (156840197) | I386G | 1 | -9.3 | -45.5 |
| 9 | 2-(2-Phenylmethoxypropan-2-yl)naphthalene-1,4-dione (139943538) | I386G | 2 | -9.3 | -37.6 |
| 10 | 2-[Hydroxy(phenyl)methyl]naphthalene-1,4-dione(245925) | I386G | 2 | -9.3 | -46.6 |

The top 10 pairings were categorized into three types based on the orientation of the underlying 1,4-naphthoquinone structure. Each type has a distinct appearance and we propose that different sets of hydrogen bonds may be present between the ligand and the protein. A distinguishing feature of Type 1 pairings is the placement of the ligand’s amine group in close proximity to FAD and N387 side chain, within 2.7 Å and 3.4 Å, respectively. Type 2 pairs are characterized by a possible hydrogen bond between the K383 residue and the ligand with an approximate length of 3.7 Å. The Type 3 pairing is similar in its orientation to Type 1; however, its ligand lacks an amine group. Still, the distance between FAD and the ligand is 4.3 Å, raising a possibility of weak electrostatic interaction. All three types may also allow hydrogen bonding with Q321 residue, with minimum residue-ligand distances ranging from 2.9 Å to 3.6 Å. All ligand atoms involved in those possible hydrogen bonds were in close proximity to electron acceptor features identified by the pharmacophore modeling of the ligands using Schrödinger Maesto [27]. Figure 4 shows the three types of pairings in greater detail.

**Figure 4:**
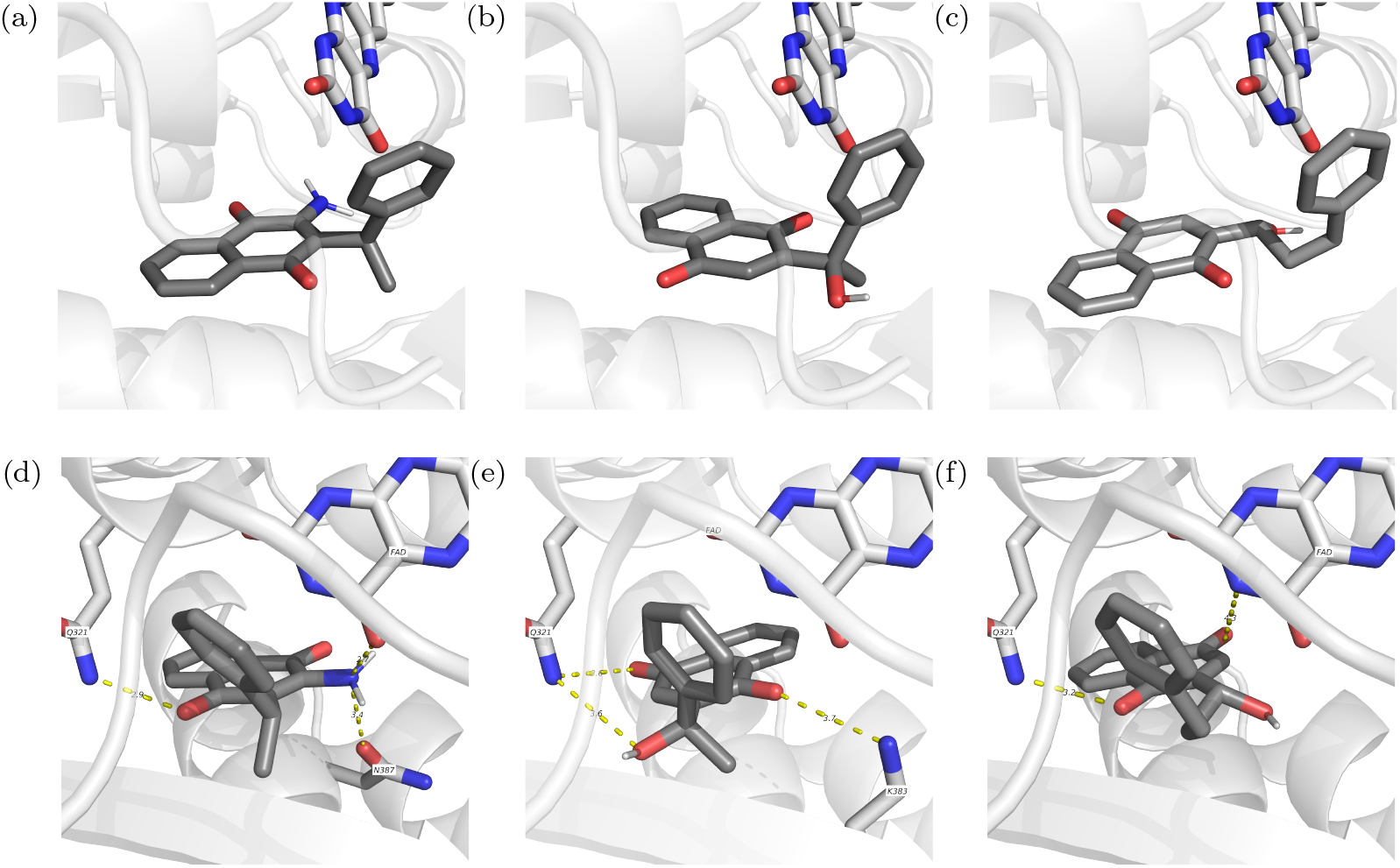
Three types of pairings observed among the top 10 co-designed protein-ligand pairs. (a-c): Type 1, 2, and 3 ligand poses seen from the same perspective, showing their relative orientation. (d-f): magnified view showing possible interactions involved in each of the Types 1, 2, and 3, respectively. The potential hydrogen bonds are shown with yellow dashed lines and numbers indicating the distance.

**Figure 5:**
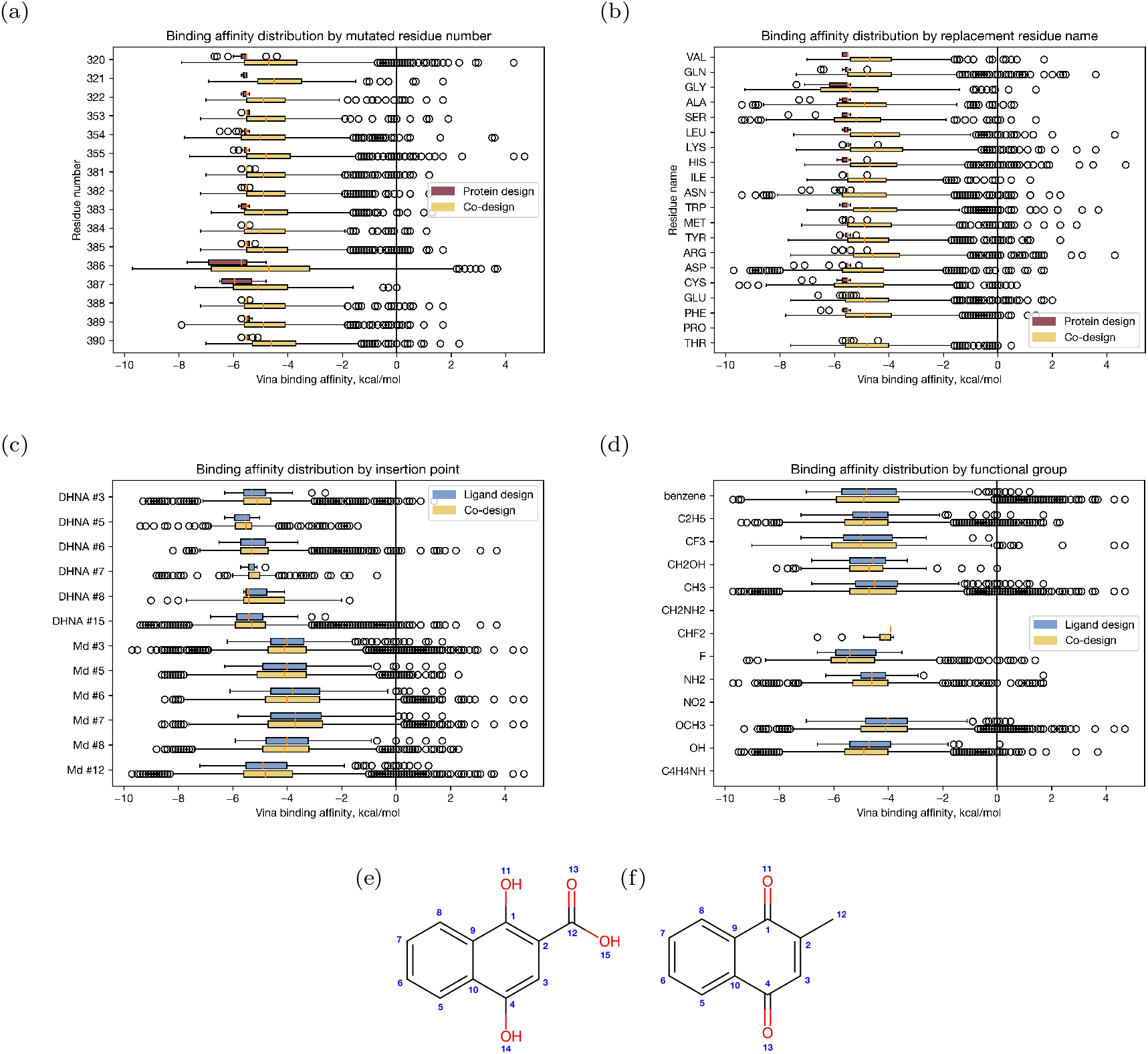
Distributions of binding affinities for individual and co-design strategies, broken down by (a) mutated residue number, (b) replacement residue name, (c) addition location on the base quinone, and (d) added functional group. Atom numbering scheme is given for (e) DHNA and (e) menadione.

Mutation of the I386 residue is a common feature of the most favorable pairings found by both the co-design and individual protein design strategies, suggesting it’s an important contributor to successful binding. This residue is part of the structure making up the quinone binding pocket and is in close contact with the ligand. In the wildtype Ndh2 sequence, this position is occupied by isoleucine, a large neutral amino acid. In mutants with improved affinity, this residue is commonly replaced by aspartic acid or a neutral polar amino acid (cysteine, serine, or asparagine). Such mutations replace a hydrophobic amino acid with a hydrophilic one. A handful of mutants replace this residue with alanine or glycine: this change has little effect on hydropathy [53], but it moves the interaction further away from the ligand as the alanine and glycine side chains are significantly smaller than that of isoleucine.

As such, it appears likely that hydropathy at this location within the binding pocket is a key factor for affinity.

The distribution of binding affinities across all sampled co-designed pairings reveals several trends. On the protein side, binding affinity was most improved by the mutation of residue 386 (Figure 5a) and the use of aspartic acid or cysteine as replacement residues (Figure 5b). On the ligand side, the largest improvement was obtained by adding functional groups at positions 5 and 15 on DHNA and 3 and 12 on menadione (Figure 5c); however, no clear trend could be seen with respect to specific functional groups (Figure 5d). Nevertheless, co-design resulted in improved binding affinities over individual design under all types of protein and ligand modifications.

## 4 Conclusion

We presented in this paper a novel paradigm for the co-design of protein-ligand pairs and its application to enhancing EET in *L. plantarum*. In contrast to traditional paradigms intended for individual design, co-design considers a larger number of protein-ligand pairings, which leads to improved binding affinities. The identified pairings were further validated using molecular dynamics calculations, pharmacophore modeling, and empirical review.

Our initial implementation of the co-design paradigm can be further developed in a number of ways. Instead of performing one round of the design cycle, it would be advantageous to carry out multiple design iterations and incorporate feedback from previous stages, transforming co-design into a comprehensive co-optimization strategy. Further, emerging methods for molecular [54, 55] and protein generation [56, 57] enable the creation of enhanced structures using ML techniques. These techniques can be further developed to more efficiently explore the protein-ligand pairings design space. Despite being in its early stages, our results highlight the crucial advantages and importance of the co-design paradigm.

## 5 Acknowledgements

This work was sponsored by Army Research Office, MURI program, contract DOD ARO #W911NF2210239.

## References

[1] Yajie Wang, Pu Xue, Mingfeng Cao, Tianhao Yu, Stephan T. Lane, and Huimin Zhao. Directed evolution: Methodologies and applications. Chemical Reviews, 121(20):12384–12444, Oct 2021.

[2] H. W. Hellinga. Rational protein design: Combining theory and experiment. Proceedings of the National Academy of Sciences, 94(19):10015–10017, 1997.

[3] Ivan V. Korendovych, Uwe T. Bornscheuer, and Matthias Höhne. Rational and Semirational Protein Design, pages 15–23. Springer New York, New York, NY, 2018.

[4] Ana Paula de Abreu, Frederico Chaves Carvalho, Diego Mariano, Luana Luiza Bastos, Juliana Rodrigues Pereira Silva, Leandro Morais de Oliveira, Raquel C. de Melo-Minardi, and Adriano de Paula Sabino. An approach for engineering peptides for competitive inhibition of the sars-cov-2 spike protein. Molecules, 29(7), 2024.

[5] Luis P.B. Scott, Jorge Chahine, and José R. Ruggiero. Using genetic algorithm to design protein sequence. Applied Mathematics and Computation, 200(1):1–9, 2008.

[6] Qianzhen Shao, Yaoyukun Jiang, and Zhongyue J. Yang. EnzyHTP computational directed evolution with adaptive resource allocation. Journal of Chemical Information and Modeling, 63(17):5650–5659, Sep 2023.

[7] Kevin K. Yang, Zachary Wu, and Frances H. Arnold. Machine-learning-guided directed evolution for protein engineering. Nature Methods, 16(8):687–694, Aug 2019.

[8] David Belanger, Suhani Vora, Zelda Mariet, Ramya Deshpande, David Dohan, Christof Angermueller, Kevin Murphy, Olivier Chapelle, and Lucy Colwell. Biological sequences design using batched bayesian optimization. In NeurIPS workshop on Bayesian Deep Learning, 2019.

[9] Johannes Linder and Georg Seelig. Fast activation maximization for molecular sequence design. BMC Bioinformatics, 22(1), October 2021.

[10] Shu-Feng Zhou and Wei-Zhu Zhong. Drug design and discovery: Principles and applications. Molecules, 22(2), 2017.

[11] Wenhao Gao and Connor W. Coley. The synthesizability of molecules proposed by generative models. Journal of Chemical Information and Modeling, 60(12):5714–5723, 2020. PMID: 32250616.

[12] Yidan Tang, Rocco Moretti, and Jens Meiler. Recent advances in automated structure-based de novo drug design. Journal of Chemical Information and Modeling, 64(6):1794–1805, Mar 2024.

[13] José Jiménez-Luna, Francesca Grisoni, and Gisbert Schneider. Drug discovery with explainable artificial intelligence. Nature Machine Intelligence, 2(10):573–584, Oct 2020.

[14] Mahesh M. Shanbhag, G. Manasa, Ronald J. Mascarenhas, Kunal Mondal, and Nagaraj P. Shetti. Fundamentals of bio-electrochemical sensing. Chemical Engineering Journal Advances, 16:100516, 2023.

[15] James U Bowie, Saken Sherkhanov, Tyler P Korman, Meaghan A Valliere, Paul H Opgenorth, and Hongjiang Liu. Synthetic biochemistry: The bio-inspired cell-free approach to commodity chemical production. Trends Biotechnol., 38(7):766–778, July 2020.

[16] James N. Blaza, Hannah R. Bridges, David Aragão, Elyse A. Dunn, Adam Heikal, Gregory M. Cook, Yoshio Nakatani, and Judy Hirst. The mechanism of catalysis by type-II NADH:quinone oxidoreductases. Scientific Reports, 7(1):40165, Jan 2017.

[17] Bruno C. Marreiros, Filipa V. Sena, Filipe M. Sousa, Ana P. Batista, and Manuela M. Pereira. Type II NADH:quinone oxidoreductase family: phylogenetic distribution, structural diversity and evolutionary divergences. Environmental Microbiology, 18(12):4697–4709, 2016.

[18] Byoung Boo Seo, Tomomi Kitajima-Ihara, Edward K. L. Chan, Immo E. Scheffler, Akemi Matsuno-Yagi, and Takao Yagi. Molecular remedy of complex I defects: Rotenone-insensitive internal NADH-quinone oxidoreductase of Saccharomyces cerevisiae mitochondria restores the NADH oxidase activity of complex I-deficient mammalian cells. Proceedings of the National Academy of Sciences, 95(16):9167–9171, 1998.

[19] Gilles Peltier, Eva-Mari Aro, and Toshiharu Shikanai. NDH-1 and NDH-2 plastoquinone reductases in oxygenic photosynthesis. Annual Review of Plant Biology, 67(1):55–80, 2016. PMID: 26735062.

[20] Sheldon W May. Applications of oxidoreductases. Current Opinion in Biotechnology, 10(4):370–375, 1999.

[21] Eric T. Stevens, Wannes Van Beeck, Benjamin Blackburn, Sara Tejedor-Sanz, Alycia R. M. Rasmussen, Emily Mevers, Caroline M. Ajo-Franklin, and Maria L. Marco. Lactiplantibacillus plantarum uses ecologically relevant, exogenous quinones for extracellular electron transfer. bioRxiv, 2023.

[22] Siliang Li, Caroline De Groote Tavares, Joe G. Tolar, and Caroline M. Ajo-Franklin. Selective bioelectronic sensing of quinone pharmaceuticals using extracellular electron transfer in Lactiplantibacillus plantarum. bioRxiv, 2023.

[23] Benjamin T. Blackburn, Robyn A. C. Alba, Vladimir O. Porokhin, Arden Hatch, Soha Hassoun, Caroline M. Ajo-Franklin, and Emily Mevers. Identifying key properties that drive redox mediator activity in Lactiplantibacillus plantarum. ChemRxiv, Manuscript submitted for publication.

[24] Oleg Trott and Arthur J. Olson. AutoDock Vina: Improving the speed and accuracy of docking with a new scoring function, efficient optimization, and multithreading. Journal of Computational Chemistry, 31(2):455–461, 2010.

[25] Stefano Forli, Ruth Huey, Michael E. Pique, Michel F. Sanner, David S. Goodsell, and Arthur J. Olson. Computational protein–ligand docking and virtual drug screening with the autodock suite. Nature Protocols, 11(5):905–919, May 2016.

[26] Lucianna H. S. Santos, Rafaela S. Ferreira, and Ernesto R. Caffarena. Integrating Molecular Docking and Molecular Dynamics Simulations, pages 13–34. Springer New York, New York, NY, 2019.

[27] Schrödinger, LLC. Schrödinger: Maestro. Release 2023-2, February 2023.

[28] Matthew P. Jacobson, David L. Pincus, Chaya S. Rapp, Tyler J.F. Day, Barry Honig, David E. Shaw, and Richard A. Friesner. A hierarchical approach to all-atom protein loop prediction. Proteins: Structure, Function, and Bioinformatics, 55(2):351–367, 2004.

[29] Matthew P. Jacobson, Richard A. Friesner, Zhexin Xiang, and Barry Honig. On the role of the crystal environment in determining protein side-chain conformations. Journal of Molecular Biology, 320(3):597–608, 2002.

[30] The UniProt Consortium. UniProt: the universal protein knowledgebase in 2021. Nucleic Acids Research, 49(D1):D480–D489, 11 2020.

[31] John Jumper, Richard Evans, Alexander Pritzel, Tim Green, Michael Figurnov, Olaf Ronneberger, Kathryn Tunyasuvunakool, Russ Bates, Augustin Žídek, Anna Potapenko, Alex Bridgland, Clemens Meyer, Simon A. A. Kohl, Andrew J. Ballard, Andrew Cowie, Bernardino Romera-Paredes, Stanislav Nikolov, Rishub Jain, Jonas Adler, Trevor Back, Stig Petersen, David Reiman, Ellen Clancy, Michal Zielinski, Martin Steinegger, Michalina Pacholska, Tamas Berghammer, Sebastian Bodenstein, David Silver, Oriol Vinyals, Andrew W. Senior, Koray Kavukcuoglu, Pushmeet Kohli, and Demis Hassabis. Highly accurate protein structure prediction with AlphaFold. Nature, 596(7873):583–589, Aug 2021.

[32] Mihaly Varadi, Stephen Anyango, Mandar Deshpande, Sreenath Nair, Cindy Natassia, Galabina Yordanova, David Yuan, Oana Stroe, Gemma Wood, Agata Laydon, Augustin Žídek, Tim Green, Kathryn Tunyasuvunakool, Stig Petersen, John Jumper, Ellen Clancy, Richard Green, Ankur Vora, Mira Lutfi, Michael Figurnov, Andrew Cowie, Nicole Hobbs, Pushmeet Kohli, Gerard Kleywegt, Ewan Birney, Demis Hassabis, and Sameer Velankar. AlphaFold Protein Structure Database: massively expanding the structural coverage of protein-sequence space with high-accuracy models. Nucleic Acids Research, 50(D1):D439–D444, 11 2021.

[33] David E. Kim, Dylan Chivian, and David Baker. Protein structure prediction and analysis using the Robetta server. Nucleic Acids Research, 32(suppl2):W526–W531, 07 2004.

[34] Minkyung Baek, Frank DiMaio, Ivan Anishchenko, Justas Dauparas, Sergey Ovchinnikov, Gyu Rie Lee, Jue Wang, Qian Cong, Lisa N. Kinch, R. Dustin Schaeffer, Claudia Millán, Hahnbeom Park, Carson Adams, Caleb R. Glassman, Andy DeGiovanni, Jose H. Pereira, Andria V. Rodrigues, Alberdina A. van Dijk, Ana C. Ebrecht, Diederik J. Opperman, Theo Sagmeister, Christoph Buhlheller, Tea Pavkov-Keller, Manoj K. Rathinaswamy, Udit Dalwadi, Calvin K. Yip, John E. Burke, K. Christopher Garcia, Nick V. Grishin, Paul D. Adams, Randy J. Read, and David Baker. Accurate prediction of protein structures and interactions using a three-track neural network. Science, 373(6557):871–876, 2021.

[35] Schrödinger, LLC. The PyMOL molecular graphics system. Version 2.5.4, August 2022.

[36] Yue Feng, Wenfei Li, Jian Li, Jiawei Wang, Jingpeng Ge, Duo Xu, Yanjing Liu, Kaiqi Wu, Qingyin Zeng, Jia-Wei Wu, Changlin Tian, Bing Zhou, and Maojun Yang. Structural insight into the type-II mitochondrial NADH dehydrogenases. Nature, 491(7424):478–482, Nov 2012.

[37] Christiam Camacho, George Coulouris, Vahram Avagyan, Ning Ma, Jason Papadopoulos, Kevin Bealer, and Thomas L. Madden. Blast+: architecture and applications. BMC Bioinformatics, 10(1):421, Dec 2009.

[38] Daniel A. DiRocco, Kevin Dykstra, Shane Krska, Petr Vachal, Donald V. Conway, and Matthew Tudge. Late-stage functionalization of biologically active heterocycles through photoredox catalysis. Angewandte Chemie International Edition, 53(19):4802–4806, 2014.

[39] Tim Cernak, Kevin D. Dykstra, Sriram Tyagarajan, Petr Vachal, and Shane W. Krska. The medicinal chemist’s toolbox for late stage functionalization of drug-like molecules. Chem. Soc. Rev., 45:546–576, 2016.

[40] M. Christina White and Jinpeng Zhao. Aliphatic c–h oxidations for late-stage functionalization. Journal of the American Chemical Society, 140(43):13988–14009, Oct 2018.

[41] Daylight Chemical Information Systems, Inc. SMARTS - a language for describing molecular patterns. https://www.daylight.com/dayhtml/doc/theory/theory.smarts.html. Accessed: 2024-01-11.

[42] RDKit contributors. RDKit: Open-source cheminformatics. Release 2022.09.1, October 2022.

[43] Sunghwan Kim, Jie Chen, Tiejun Cheng, Asta Gindulyte, Jia He, Siqian He, Qingliang Li, Benjamin A Shoemaker, Paul A Thiessen, Bo Yu, Leonid Zaslavsky, Jian Zhang, and Evan E Bolton. PubChem 2023 update. Nucleic Acids Research, 51(D1):D1373–D1380, 10 2022.

[44] G. Madhavi Sastry, Matvey Adzhigirey, Tyler Day, Ramakrishna Annabhimoju, and Woody Sherman. Protein and ligand preparation: parameters, protocols, and influence on virtual screening enrichments. Journal of Computer-Aided Molecular Design, 27(3):221–234, Mar 2013.

[45] Schrödinger, LLC. Ligprep. Release 2023-2, February 2023.

[46] Andrew Waterhouse, Martino Bertoni, Stefan Bienert, Gabriel Studer, Gerardo Tauriello, Rafal Gumienny, Florian T Heer, Tjaart A P de Beer, Christine Rempfer, Lorenza Bordoli, Rosalba Lepore, and Torsten Schwede. SWISS-MODEL: homology modelling of protein structures and complexes. Nucleic Acids Research, 46(W1):W296–W303, 05 2018.

[47] Stefan Bienert, Andrew Waterhouse, Tjaart A. P. de Beer, Gerardo Tauriello, Gabriel Studer, Lorenza Bordoli, and Torsten Schwede. The SWISS-MODEL Repository—new features and functionality. Nucleic Acids Research, 45(D1):D313–D319, 11 2016.

[48] Nicolas Guex, Manuel C. Peitsch, and Torsten Schwede. Automated comparative protein structure modeling with SWISS-MODEL and Swiss-PdbViewer: A historical perspective. ELECTROPHORESIS, 30(S1):S162–S173, 2009.

[49] Gabriel Studer, Christine Rempfer, Andrew M Waterhouse, Rafal Gumienny, Juergen Haas, and Torsten Schwede. QMEANDisCo—distance constraints applied on model quality estimation. Bioinformatics, 36(6):1765–1771, 11 2019.

[50] Martino Bertoni, Florian Kiefer, Marco Biasini, Lorenza Bordoli, and Torsten Schwede. Modeling protein quaternary structure of homo- and hetero-oligomers beyond binary interactions by homology. Scientific Reports, 7(1):10480, Sep 2017.

[51] Markus Wiederstein and Manfred J. Sippl. ProSA-web: interactive web service for the recognition of errors in three-dimensional structures of proteins. Nucleic Acids Research, 35(suppl2):W407–W410, 07 2007.

[52] Manfred J. Sippl. Recognition of errors in three-dimensional structures of proteins. Proteins: Structure, Function, and Bioinformatics, 17(4):355–362, 1993.

[53] Christelle Pommié, Séverine Levadoux, Robert Sabatier, Gérard Lefranc, and Marie-Paule Lefranc. Imgt standardized criteria for statistical analysis of immunoglobulin v-region amino acid properties. Journal of Molecular Recognition, 17(1):17–32, 2004.

[54] Viraj Bagal, Rishal Aggarwal, P. K. Vinod, and U. Deva Priyakumar. Molgpt: Molecular generation using a transformer-decoder model. Journal of Chemical Information and Modeling, 62(9):2064–2076, May 2022.

[55] Youjun Xu, Kangjie Lin, Shiwei Wang, Lei Wang, Chenjing Cai, Chen Song, Luhua Lai, and Jianfeng Pei. Deep learning for molecular generation. Future Medicinal Chemistry, 11(6):567–597, 2019. PMID: 30698019.

[56] Surojit Biswas, Grigory Khimulya, Ethan C. Alley, Kevin M. Esvelt, and George M. Church. Low-n protein engineering with data-efficient deep learning. Nature Methods, 18(4):389–396, Apr 2021.

[57] Ali Madani, Ben Krause, Eric R. Greene, Subu Subramanian, Benjamin P. Mohr, James M. Holton, Jose Luis Olmos, Caiming Xiong, Zachary Z. Sun, Richard Socher, James S. Fraser, and Nikhil Naik. Large language models generate functional protein sequences across diverse families. Nature Biotechnology, 41(8):1099–1106, Aug 2023.

