## Supplementary Materials for "Protein-ligand co-design: a case for improving binding affinity between Type II NADH:quinone oxidoreductase and quinones"

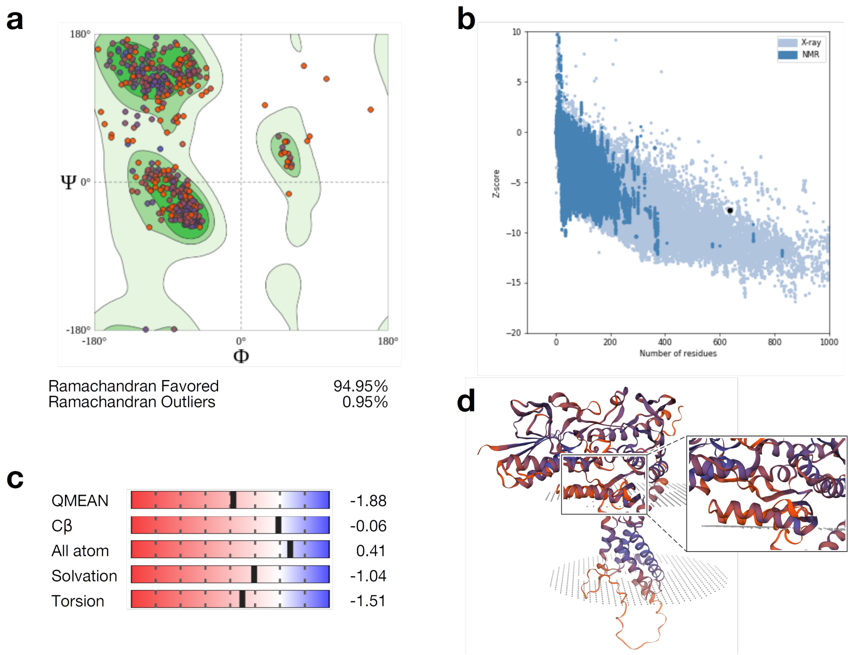


Figure S1: Structural model quality validation for the *L. plantarum* strain of Ndh2. (a) Ramachandran plot showing (Φ, Ψ) residue angles in mostly favorable regions (94.95%). (b) Pro-SA Z-score of model (-7.76, black dot) was within the range of known protein structures solved by x-ray crystallography for its sequence length. (c) QMEAN assesment scores describing model quality based on torsion angles and solvation. (d) Structural model annotated with per-residue confidence values predicted by SWISS-MODEL, the insert shows the quinone binding site magnified.

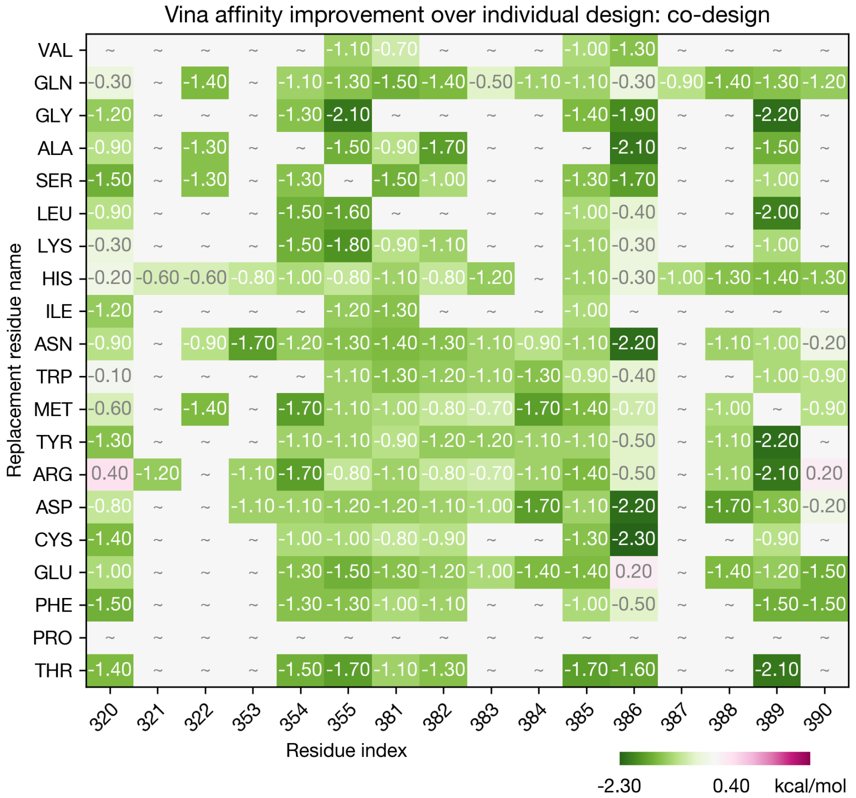


Figure S2: Maximum change in AutoDock Vina binding affinity accomplished by the co-design strategy over protein individual design, as a function of mutation site and replacement amino acid. In all cases, except for V320R, I386E, and Y390R, co-design identified more favorable pairings, with affinity changes as large as -2.5 kcal/mol. Protein variants not explored due to the replacement residue being identical to the original one, high Δ Prime from Residue Scanning, or those unable to be docked successfully, are marked by the "∼" symbol.

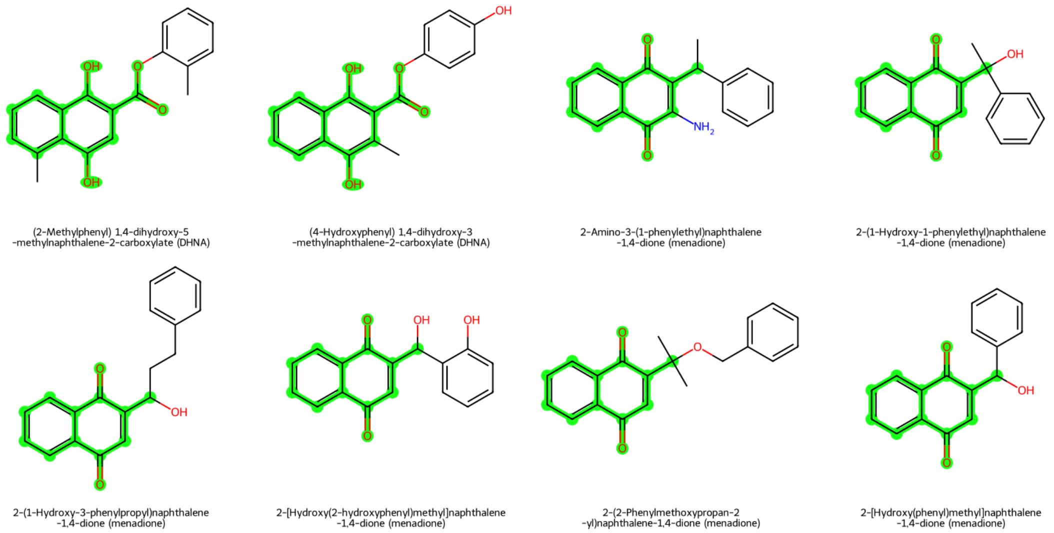


Figure S3: Ligands corresponding to the top 10 poses suggested by the co-design approach. The names of ancestor quinones are given in parentheses. The structures of ancestor quinones are highlighted in green.
